# Impedance flow cytometry for viability analysis of *Corynebacterium glutamicum*

**DOI:** 10.1101/2021.09.23.461525

**Authors:** Fabian Stefan Franz Hartmann, Ioannis Anastasiou, Tamara Weiß, Jing Shen, Gerd Michael Seibold

## Abstract

*Corynebacterium glutamicum* efficiently produces glutamate when growth is inhibited. Analyses of viability in this non-growing state requires time consuming plating and determination of colony forming units. We here establish impedance flow cytometry measurements to assess the viability of non-growing, glutamate producing *C. glutamicum* cultures within minutes.

## INTRODUCTION

Biomass and viability are key parameters in cultivation processes and different techniques have been developed for *on-line* or *in-situ* measurements of these two variables (Olsson and Nielsen, 1997; Reichelt et al., 2016). They are commonly assessed by measuring changes of the optical density (OD), cell dry weight (CDW), total cell numbers (colony forming units (CFU)), live-dead-staining using fluorescent dyes, or metabolic activity (*e.g*. offgas analyses of CO_2_ formation and O_2_ consumption, *at-line* analyses of substrate and product concentrations). Impedance flow cytometry (IFC) *via* microfluidic sensors was initially developed to accurately assess bacteria levels in drinking water in real-time (Clausen et al., 2018) and to detect pathogens (Furst and Francis, 2019). The technology was described to be able to detect all types of bacteria and to distinguish them from particles and dead cells based on their electrical properties (Clausen et al., 2018; Furst and Francis, 2019). Additionally, it is considered as a novel tool to quickly assess viability of bacteria during bench-scale cultivations as well as industrial fermentations (Bertelsen et al., 2020; David et al., 2012).

The Gram-positive bacterium *C. gutamicum* is used for large scale industrial production of amino acids (Eggeling and Bott, 2015) and has been engineered into a versatile platform organism for the production of fine and bulk chemicals from renewable feedstocks (Wolf et al., 2021). Determining the cellular fitness of *C. glutamicum* during glutamate production has limitations as glutamate secretion is triggered by conditions disturbing cell wall and/or membrane integrity (Eggeling et al., 2001; Radmacher et al., 2005). Thus, glutamate producing cultures of *C. glutamicum* consist of non-growing but optimally viable and active cells, a physiological state which cannot easily be assessed *via* simple biomass measurements.

In this communication we investigate the use of a recently developed, hand held multi-frequency IFC device for the analysis of the viability of *C. glutamicum* cultures in different physiological states. We compare IFC measurements with analyses of CFU and finally analyze the viability of *C. glutamicum* cells during penicillin induced glutamate production.

## RESULTS & DISCUSSION

For the comparison of IFC with different established techniques commonly used for analyses of *C. glutamicum* viability, the wild type strain *C. glutamicum* ATCC13032 (Abe et al., 1967) was cultivated in 50 mL 2xTY-medium at 30 °C in shake flasks. Samples of exponentially growing *C. glutamicum* cells were harvested by centrifugation, washed once with saline (0.9 g/L NaCl), and incubated for 30 min. with either saline to preserve living cells or 70% (v/v) isopropyl alcohol to obtain dead cells (Seibold et al., 2009). Subsequently, the treated cells were pelleted and then re-suspended in saline. Cell suspensions were plated on 2xTY-agar plates and incubated for 48 h at 30° C. Finally, 3.0 × 10^8^ ± 9.2 × 10^6^ CFU/mL were detected for saline incubated cells, whereas no CFU were detected when 100 μl of un-diluted suspension of isopropyl alcohol treated cells were plated (Fig. 1a). When plating mixtures of both suspensions, the CFU was determined to be higher upon adjusting a higher (60%) amount of saline-incubated cell suspension in the respective mixture, whereas it was *vice versa* when applying a lower amount (40%) (Fig. 1a). This shows that by mixing the differently pretreated cell suspension, mixtures containing a defined ratio of viable and dead cells can accurately be set.

**Figure 1:**
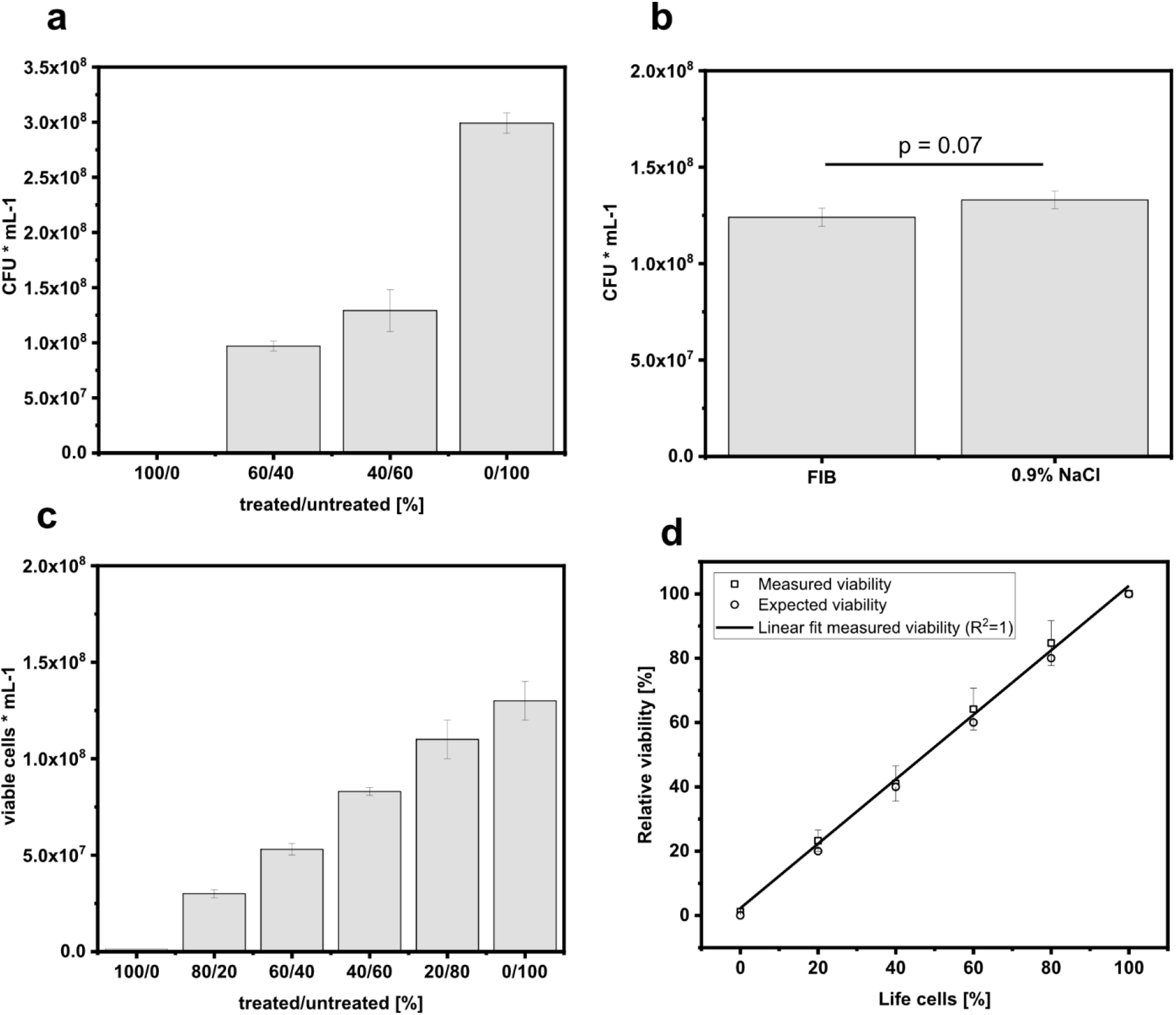
Viability measurements of *C. glutamicum* cultures *via* different methods. Colony forming units (CFU) were determined for cultures treated with isopropyl alcohol (treated) or saline (untreated) and different mixtures of both types of cultures (a). CFU numbers determined and compared for *C. glutamicum* cultures suspended in flow impedance buffer (FIB) or saline (b). Determination of viable cells *via* impedance flow cytometry (IFC) for *C. glutamicum* cultures treated with isopropyl alcohol (treated) or saline (untreated) and different mixtures of both (c). Linear regression curve of measured relative cell viability using IFC and expected relative viabilities for the different prepared culture mixtures (d). One-way ANOVA with Tukey’s test was performed to assess the significance of difference between CFU numbers for *C. glutamicum* cultures suspended in different buffers. Numbers were determined to be not significantly different (p < 0.05 is considered as significantly different). Error bars represent standard deviation from at least three replicates.

IFC measurements of bacterial viability require a low ionic strength of the buffer used to prepare the suspension (Clausen et al., 2018; David et al., 2012). Therefore, Flow Impendance Buffer (FIB) as an alternative to saline to suspend *C. glutamicum* cells was tested. Upon incubation of *C. glutamicum* cells for 30 min. with either saline or FIB (0.005% PBS in MilliQ water, conductivity below 8 mS), the CFU was determined to be 1.33 × 10^8^ ± 4.55 × 10^6^ and 1.24 × 10^8^ ± 4.68 × 10^6^, respectively, which was analyzed to be not significantly different (p = 0.07) (Fig. 1b).

When suspensions of defined amounts of dead and viable *C. glutamicum* cultures were analyzed *via* IFC, 1.3 × 10^8^ ± 1.0 × 10^7^ viable cells/mL were detected for non-treated cells (Fig. 1c), which is in the same range of the number of CFU determined for FIB suspensions as well as saline diluted cells (see above). The number of viable cells detected *via* IFC in the suspension decreased when the mixtures contained isopropyl alcohol treated cells (Fig. 1c). Indeed, a linear correlation (R^2^=1) between the percentage of living cells in the defined suspensions and the relative viability measured *via* IFC was determined (Fig. 1d). The results indicate that IFC is well suited to analyze viability of *C. glutamicum* cells in suspensions containing both viable and non-viable cells.

The addition of penicillin inhibits growth of *C. glutamicum* and in turn triggers glutamate production (Eggeling et al., 2001). Effects of penicillin on growth of *C. glutamicum* in CGXII minimal medium (Graf et al., 2018) with 1 % (w/v) glucose as sole carbon source were tested in a BioLectorII system (m2p-labs, Baesweiler/DE) using 48-well Flowerplates (m2p-labs, Baesweiler/DE) and 800 μL cultures at 1500 rpm and 30°C as recently described (Hemmerich et al., 2019). As depicted in Fig. 2a, varying the penicillin concentration between 0 and 1000 U/mL affected growth and final biomass of *C. glutamicum*. Growth of *C. glutamicum* ceased from a growth rate of 0.17 ± 0.01 h^−1^ in absence of penicillin to a growth rate of 0.01 ± 0.01 h^−1^ for concentrations above 1 U/mL (Fig. 2b). To confirm glutamate production by non-growing *C. glutamicum* strains, a glutamate assay was performed according to the manufacturer’s instructions (Megazyme, Ireland). As expected, glutamate production was decoupled from growth, resulting in a final titer of 0.03 ± 0 g/L glutamate in absence or presence of low penicillin concentrations, but increased in presence of > 1 U/mL penicillin (Fig. 2 b) to titers of up to 0.47 ± 0.01 g/L after 24 h of cultivation. The formation of glutamate by the cultures in presence of more than 1 U/mL penicillin indicates that despite the fact that no biomass formation was observed, active cells were present. Indeed, the viability measurements *via* plating and CFU determination (Fig 2c) as well as *via* IFC (Fig. 2d) confirmed that even at a concentration of 1000 U/mL penicillin, viable cells are present in the *C. glutamicum* cultures. In detail for cells treated with 0.1 U/ mL penicillin 2.88 × 10^7^ ± 2.9 × 10^6^ CFU * mL^−1^ * OD_600nm_^−1^ and 1.31 ×10^8^ ± 7.66 × 10^6^ viable cells * mL^−1^ * OD_600nm_^−1^ were determined *via* plating and IFC, respectively. This corresponds to 46 ± 15 % and 42 ± 5 % of the total cell numbers determined for non-penicillin treated cultures *via* plating and IFC, respectively. With higher penicillin concentration the relative cell viability determined *via* the two different methods remained approximately between 30% - 45% for most approaches. This fits to the observation that in presence of high penicillin concentrations cells were in a non-growing but producing state. To note, higher absolute cell numbers were determined *via* IFC when compared to CFU (plating) (Fig. 2c, d). This observation might indicate that cultures under these conditions contained viable but not culturable cells (Oliver, 2005), as it has been reported for *C. glutamicum* cadaverine-production strains (Olughu et al., 2020).

**Figure 2:**
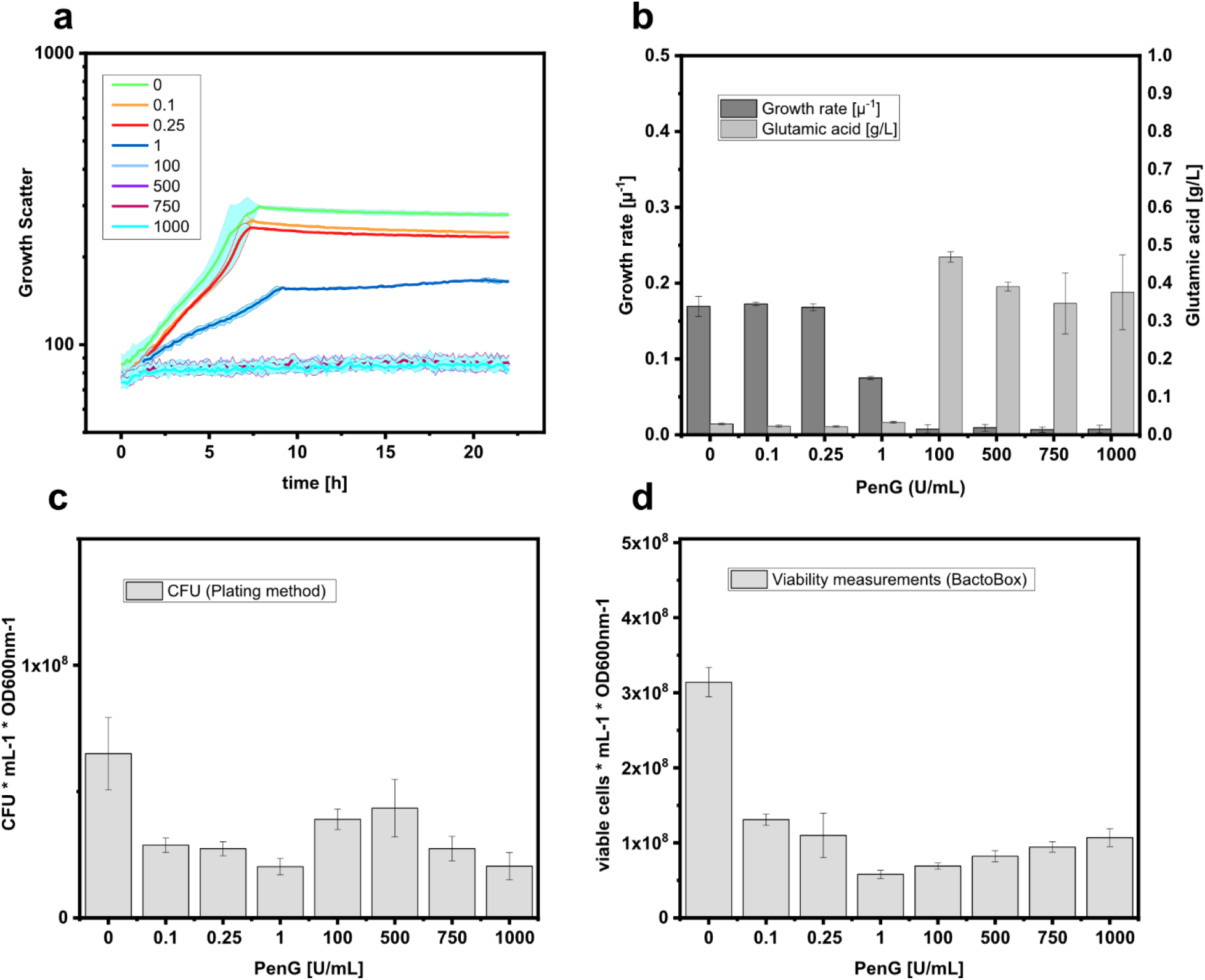
Growth of *C. glutamicum* in minimal medium (CGXII) at different set penicillin concentrations (0-1000 U/mL) (a), growth rates and final glutamate titers in the cultures supernatant when growing in presence of different penicillin concentrations (b) and the final cell viability determined *via* plating (c) and by impedance flow cytometry (IFC) using the BactoBox (d). Cultivation was performed in a BioLectorII system (m2p-labs, Baesweiler/DE) using 48-well Flowerplates (m2p-labs, Baesweiler/DE) and 800 μL cultures at 1500 rpm and 30°C with 1% (w/v) glucose as sole carbon source. Error bars represent standard deviation from biological triplicates.

Taken together, the results show that IFC allows fast analyses of the cellular viability of *C. glutamicum*. The procedure, including sampling and analysis, takes no longer than five minutes to determine the viability of a cell culture. This is crucial for fast decision-making for processes involving non-growing but producing strains.

## AUTHOR’S CONTRIBUTION

**Fabian Stefan Franz Hartmann:** Conceptualization, Investigation, Data curation, Methodology, Visualization, Formal analysis, Validation, Writing- Original draft, Review & Editing

**Ioannis Anastasiou:** Investigation, Data curation, Formal analysis, Validation

**Tamara Weiß:** Investigation, Methodology, Writing-Review & Editing

**Jing Shen:** Methodology, Writing- Review & Editing

**Gerd Michael Seibold:** Conceptualization, Methodology, Supervision, Writing- Original draft, Review & Editing

## ACKNOWLEDGEMENTS

We would like to thank the Fermentation Core at DTU Bioengineering for excellent technical support and Gustav E. Skands and Michael Maimann of SBT Instruments A/S for very helpful discussions.

